## Supplemental Materials for "A nanopore interface for high bandwidth DNA computing"

Tables S1-S4

Figures S1-S7

| Circuit # | Circuit Component | Sequence | Purification Method |
|---|---|---|---|
| 0 | Gate | AGAGTGTGGAGTTGATAGGAGAG | Standard Desalting |
| | Output | ACACCTTACTCTCTACTCTCCTATCAACTCCACATTTTTTNNNNNNTT/3BioTEG/ | HPLC |
| | Input | CCTATCAACTCCACACTCTCACTAATTCTACATC | Standard Desalting |
| | Fuel | TACCTCATTCAAACTCTCTCCTATCAACTCCACA | Standard Desalting |
| | Quencher | /5IAbRQ/ACACCTTACTCTCTA | HPLC |
| | Fluorophore | AGAGTAGAGAGTAAGGTGT/36-FAM/ | HPLC |
| 1 | Gate | AGAGAATAATGGTTGTAGGAGAG | Standard Desalting |
| | Output | CTTCTTATACCACACCTCTCCTACAACCATTATTTTTTTTNNNNNNCA/3BioTEG/ | HPLC |
| | Input | CCTACAACCATTATTCTCTCAATCTACCAAACTC | Standard Desalting |
| | Fuel | ACAAATACCTCATCCCTCTCCTACAACCATTATT | Standard Desalting |
| | Quencher | /5IAbRQ/CTTCTTATACCACAC | HPLC |
| | Fluorophore | AGAGGTGTGGTATAAGAAG/36-FAM/ | HPLC |
| 2 | Gate | AGAGTAAGTATAGAGGTGAAGAG | Standard Desalting |
| | Output | CAACTACAATCTCTCCTCTTCACCTCTATACTTATTTTTTNNNNNNTA/3BioTEG/ | HPLC |
| | Input | TCACCTCTATACTTACTCTTCCTTCATCTTCTAC | Standard Desalting |
| | Fuel | TCTCCAATTTCAACTCTCTTCACCTCTATACTTA | Standard Desalting |

| | | | |
|---|---|---|---|
| | Quencher | /5IAbRQ/CAACTACAATCTCTC | HPLC |
| | Fluorophore | AGAGGAGAGATTGTAGTTG/36-FAM/ | HPLC |
| 3 | Gate | AGAGAGGATTAGGATAGTGAGAG | Standard Desalting |
| | Output | AACCACATTAACCTTCTCTCACTATCCTAATCCTTTTTTTNNNNNNCC/3BioTEG/ | HPLC |
| | Input | CACTATCCTAATCCTCTCTAAACCTTACCACCAC | Standard Desalting |
| | Fuel | CCTTCTCAACTCCTCCTCTCACTATCCTAATCCT | Standard Desalting |
| | Quencher | /5IAbRQ/AACCACATTAACCTT | HPLC |
| | Fluorophore | AGAGAAGGTTAATGTGGTT/36-FAM/ | HPLC |
| 4 | Gate | AGAGGGTGTTTAGAGTTTAAGAG | Standard Desalting |
| | Output | TTATCCAACTCACTACTCTTAAACTCTAAACACCTTTTTTNNNNNNAT/3BioTEG/ | HPLC |
| | Input | TAAACTCTAAACACCCTCTAATAACACCTCCTAA | Standard Desalting |
| | Fuel | CTCTTCTTTCCAAACCTCTTAAACTCTAAACACC | Standard Desalting |
| | Quencher | /5IAbRQ/TTATCCAACTCACTA | HPLC |
| | Fluorophore | AGAGTAGTGAGTTGGATAA/36-FAM/ | HPLC |
| 5 | Gate | AGAGAAAGTGATAAGATGGAGAG | Standard Desalting |
| | Output | TCTTTCACCTCACATCTCTCCATCTTATCACTTTTTTTTTNNNNNNCA/3BioTEG/ | HPLC |
| | Input | CCATCTTATCACTTTCTCTATTACTTCCTACACC | Standard Desalting |

| | | | |
|---|---|---|---|
| | Fuel | ATCCTCCTTCCATCCCTCTCCATCTTATCACTT T | Standard Desalting |
| | Quencher | /5IAbRQ/TCTTTCACCTCACAT | HPLC |
| | Fluorophore | AGAGATGTGAGGTGAAAGA/3Cy5Sp/ | HPLC |
| 6 | Gate | AGAGGGTATTAGTTAGGTAAGAG | Standard Desalting |
| | Output | TCCATTTCATTTCACCTCTTACCTAACTAATAC CTTTTTTNNNNNNAA/3BioTEG/ | HPLC |
| | Input | TACCTAACTAATACCCTCTCTCCATAACATTC CA | Standard Desalting |
| | Fuel | ACTTCTAACAACTACCTCTTACCTAACTAATA CC | Standard Desalting |
| | Quencher | /5IAbRQ/TCCATTTCATTTCAC | HPLC |
| | Fluorophore | AGAGGTGAAATGAAATGGA/36-FAM/ | HPLC |
| 7 | Gate | AGAGTGTTAGTAGTAGAGTAGAG | Standard Desalting |
| | Output | AAATTCTATCCACTCCTCTACTCTACTACTAA CATTTTTTNNNNNNCC/3BioTEG/ | HPLC |
| | Input | ACTCTACTACTAACACTCTTCTACATCCACAT CT | Standard Desalting |
| | Fuel | CACTCAATAACTACCCTCTACTCTACTACTAA CA | Standard Desalting |
| | Quencher | /5IAbRQ/AAATTCTATCCACTC | HPLC |
| | Fluorophore | AGAGGAGTGGATAGAATTT/36-FAM/ | HPLC |
| 8 | Gate | AGAGGTATAAAGGAGTTTGAGAG | Standard Desalting |
| | Output | TAACTCTACCACAAACTCTCAAACTCCTTTAT ACTTTTTTNNNNNNTA/3BioTEG/ | HPLC |

| | | | |
|---|---|---|---|
| | Input | CAAACTCCTTTATACCTCTCTCTACTCATCTTC C | Standard Desalting |
| | Fuel | CCACCTCCATCTATACTCTCAAACTCCTTTAT AC | Standard Desalting |
| | Quencher | /5IAbRQ/TAACTCTACCACAAA | HPLC |
| | Fluorophore | AGAGTTTGTGGTAGAGTTA/3Cy5Sp/ | HPLC |
| 9 | Gate | AGAGTGGTAAGGTAGTTAAAGAG | Standard Desalting |
| | Output | CTAACAAACTTTACCCTCTTTAACTACCTTAC CATTTTTTNNNNNNTC/3BioTEG/ | HPLC |
| | Input | TTAACTACCTTACCACTCTACATTCCTTCTAAT C | Standard Desalting |
| | Fuel | ATTTACATCTCAACCCTCTTTAACTACCTTAC CA | Standard Desalting |
| | Quencher | /5IAbRQ/CTAACAAACTTTACC | HPLC |
| | Fluorophore | AGAGGGTAAAGTTTGTTAG/3Cy5Sp/ | HPLC |

**Table S1:** Table of seesaw gate, output strand, input strand, fuel strand, quencher-labeled strand, and fluorophore-labeled strand sequences for all ten circuits used in kinetics analysis and multiplexing experiments. Iowa Black RQ from IDT is used as the quencher molecule and either 6-FAM or Cy 5 is used as the fluorophore in the fluorescent reporter complexes. The nanopore barcode region (red) can be substituted with any barcode from **Table S2**.

| Barcode # | Set A | Set B | Set C |
|---|---|---|---|
| 0 | CAAATA | GGGTTC | /iSpC3/CATAC |
| 1 | TCATAC | TGATTG | T/iSpC3/ATAC |
| 2 | ATATCT | AGAGTT | TC/iSpC3/TAC |
| 3 | CTCCAC | AGAGGA | TCA/iSpC3/AC |
| 4 | ATCTAA | ATATCA | TCAT/iSpC3/C |
| 5 | CTCAAA | TTCTGT | TCATA/iSpC3/ |
| 6 | AAATAC | AGCCTC | /iSpC3/CATA/iSpC3/ |
| 7 | TCCAAC | GATACT | T/iSpC3/AT/iSpC3/C |
| 8 | CAAAAC | TCTCTG | TC/iSpC3//iSpC3/AC |
| 9 | ACCTCC | AATCAA | T/iSpC3//iSpC3/TAC |
| 10 | - | TGGAAG | TCA/iSpC3//iSpC3/C |
| 11 | - | GCACAT | TCAT/iSpC3//iSpC3/ |
| 12 | - | - | T/iSpC3//iSpC3//iSpC3/AC |
| 13 | - | - | AA/iSpC3/CAA |

**Table S2:** Table of all explored output strand barcodes organized into Set A (randomly selected), Set B (based on predictive model) and Set C (contains abasic sites). Abasic sites are denoted as /iSpC3/ (C3 Spacer phosphoramidite modification from IDT) .

| **Circuit Component** | **Sequence** | **Purification Method** |
| --- | --- | --- |
| Gate | TGAGTGTGATTGTGTTATGAGTG | Standard Desalting |
| Output | CAACATATCAATTCACTCATAACACAATCACATTTTTT TCATACCA/3BioTEG/ | HPLC |
| Input | CATAACACAATCACACTCACCACCAAACTTCA | Standard Desalting |
| Fuel | CACTAACATACAACACTCATAACACAATCACA | Standard Desalting |
| Quencher | /5IAbRQ/CAACATATCAATTCA | HPLC |
| Fluorophore | TGAGTGAATTGATATGTTG/3Cy5Sp/ | HPLC |

**Table S3:** Table of seesaw gate, output strand, input strand, fuel strand, quencher-labeled strand, and fluorophore-labeled strand sequences for clamped circuit from **Figure S1**. Iowa Black RQ from IDT is used as the quencher molecule and Cy 5 is used as the fluorophore in the reporter complex. Barcode A1 (red) from **Table S2** was used in this circuit's output strand.

| **Probe Component** | **Sequence** | **Purification Method** |
|---|---|---|
| **let-7a Probe** | | |
| Input | UGAGGUAGUAGGUUGUAUAGUU | HPLC |
| Helper | GAGTGTAGTGGAAGTTGGAG | PAGE |
| Bottom | AACTATACAACCTACTACCTCACTCCAACTTCCACTACACTC/36-FAM/ | HPLC |
| Top | /5IAbRQ/GAGTGTAGTGGAAGT | HPLC |
| Output | TGGAGTGAGGTAGTAGGTTTTTTTTTGGAAGTT/3BioTEG/ | HPLC |
| **let-7c Probe** | | |
| Input | UGAGGUAGUAGGUUGUAUGG | HPLC |
| Helper | TGAGGTATGGAGTGAGTGGA | PAGE |
| Bottom | AACCATACAACCTACTACCTCATCCACTCACTCCATACCTCA/36-FAM/ | HPLC |
| Top | /5IAbRQ/TGAGGTATGGAGTGA | HPLC |
| Output | GTGGATGAGGTAGTAGGTTTTTTTTAA/iSpC3/CAATT/3BioTEG/ | HPLC |
| **let-7e Probe** | | |
| Input | UGAGGUAGGAGGUUGUAUAGUU | HPLC |
| Helper | TGAGGTATGGAGTGAGTGGA | PAGE |
| Bottom | AACTATACAACCTCCTACCTCATCCACTCACTCCATACCTCA/3Cy5Sp/ | HPLC |
| Top | /5IAbRQ/TGAGGTATGGAGTGA | HPLC |
| Output | GTGGATGAGGTAGGAGGTTTTTTTTT/iSpC3//iSpC3//iSpC3/ACTT/3BioTEG/ | HPLC |

**Table S4:** Table of let-7 miRNA input, helper strand, bottom strand (with fluorophore), top strand (with quencher), and output strand (with barcode) for the two-step let-7 detection probes. The barcode region is represented in red. Barcodes B10, C13, and C12 from Table S2 were used for the let-7a, let-7c, and let-7e probes respectively.

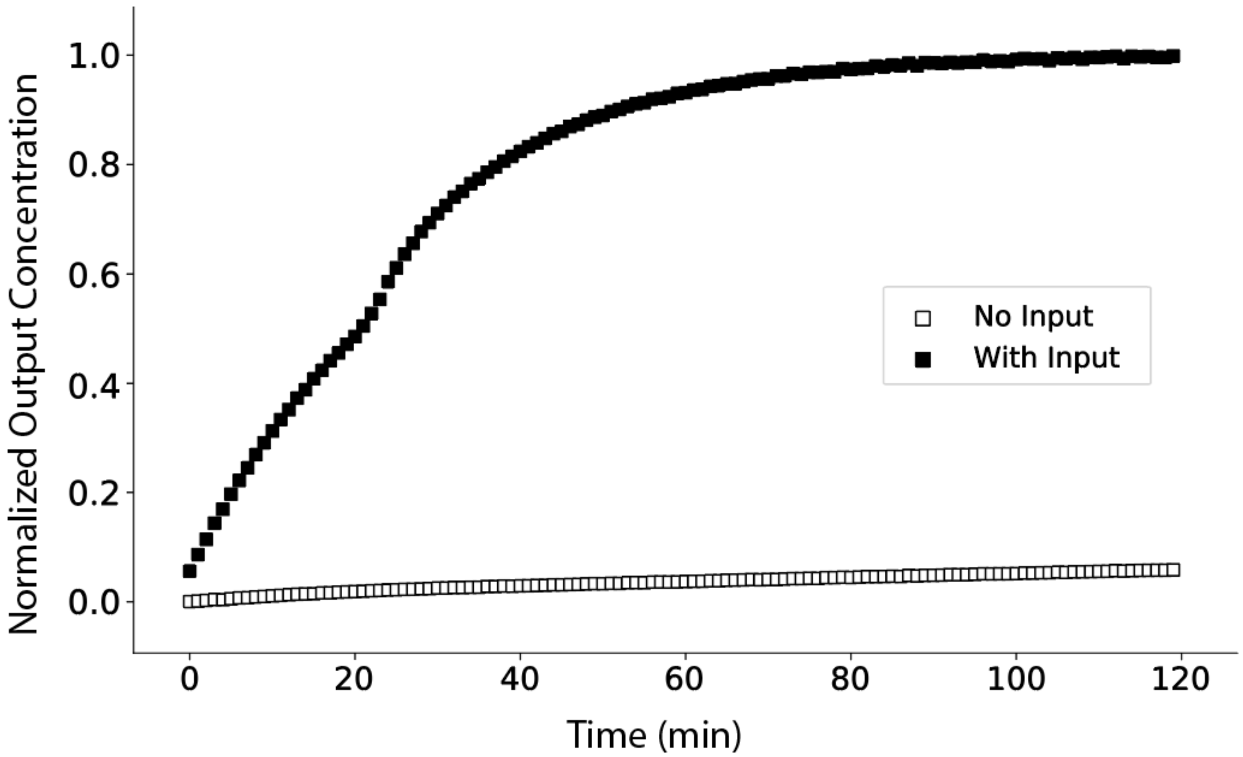


**Figure S1:** Kinetics of a clamped catalytic DSD circuit measured on a fluorospectrometer. Addition of clamp domains mitigate circuit leakage caused by blunt end stacking.

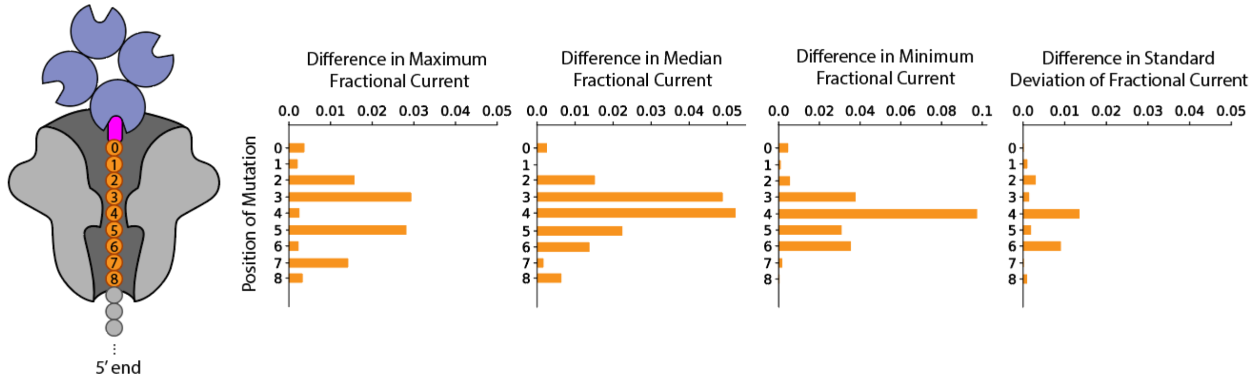


**Figure S2:** Each plot depicts the change in either maximum, median, minimum, or standard deviation of nanopore fractional current elicited by a single-nucleotide mutation at each position on the strand's barcode.

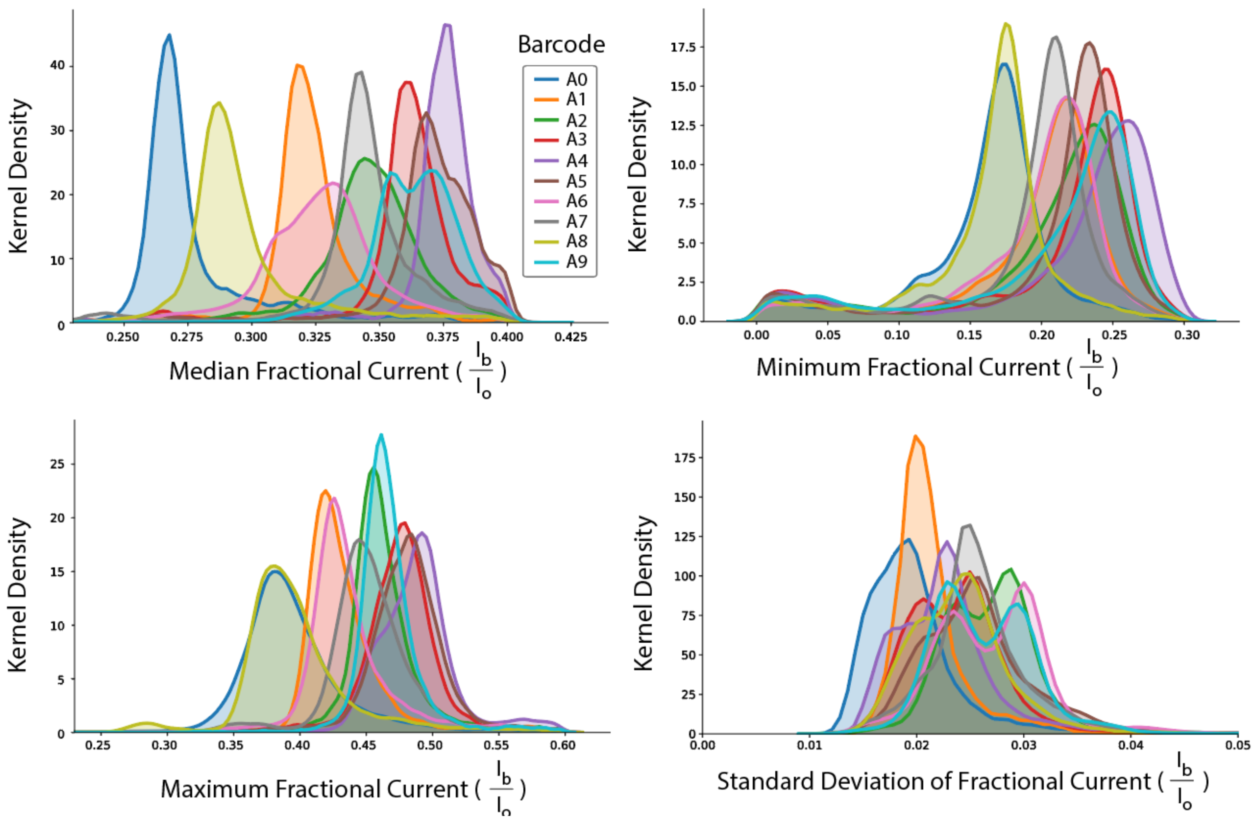


**Figure S3:** Distributions of median, minimum, maximum, and standard deviation of nanopore fractional current for each barcode in Set A. Each distribution is composed of ~14500 data points.

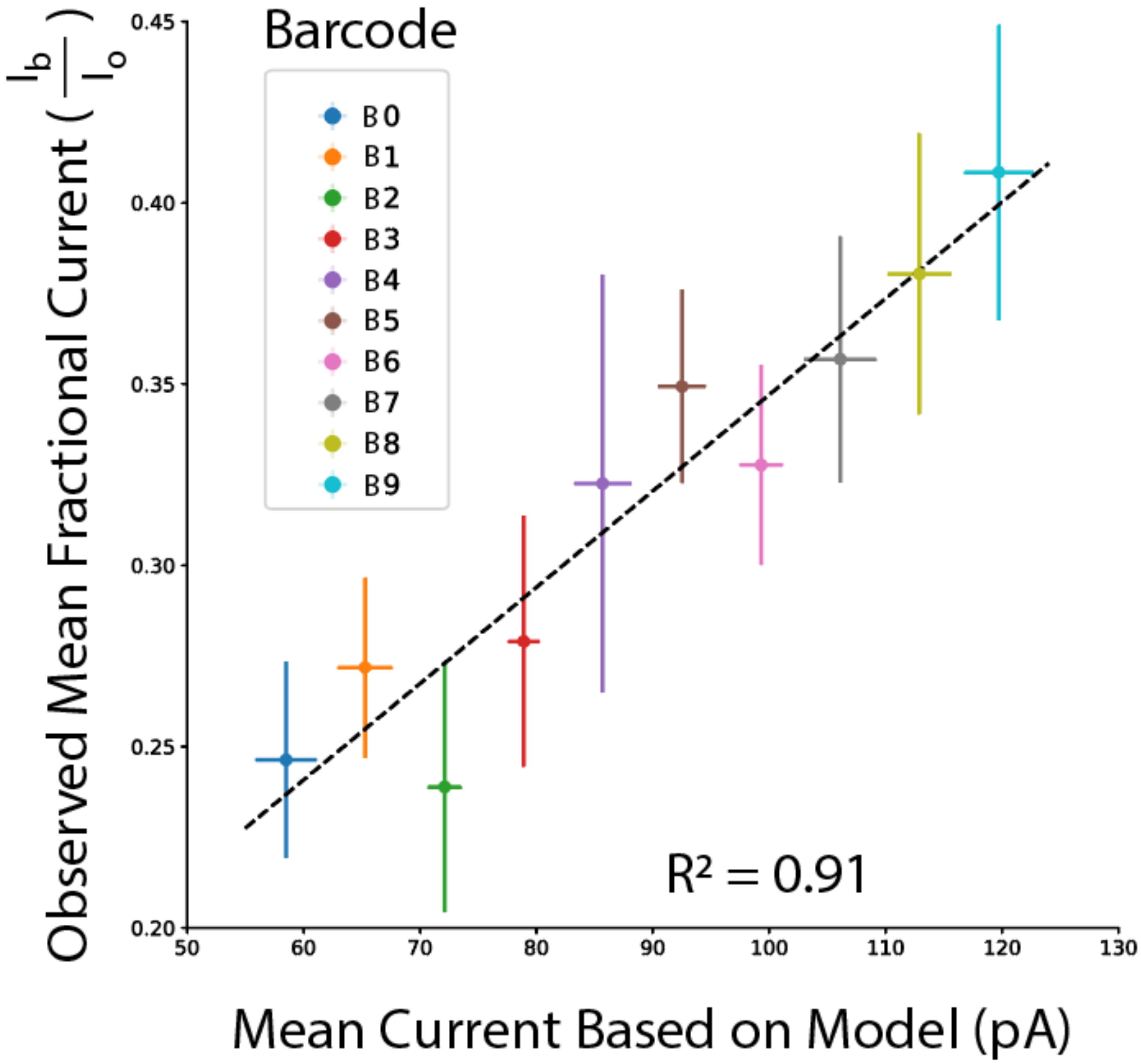


**Figure S4:** Correlation of observed mean fractional currents to the mean current from predictive model for barcodes B1-B9. Vertical and horizontal error bars show standard deviation for observed mean fractional current and mean current based on model respectively.

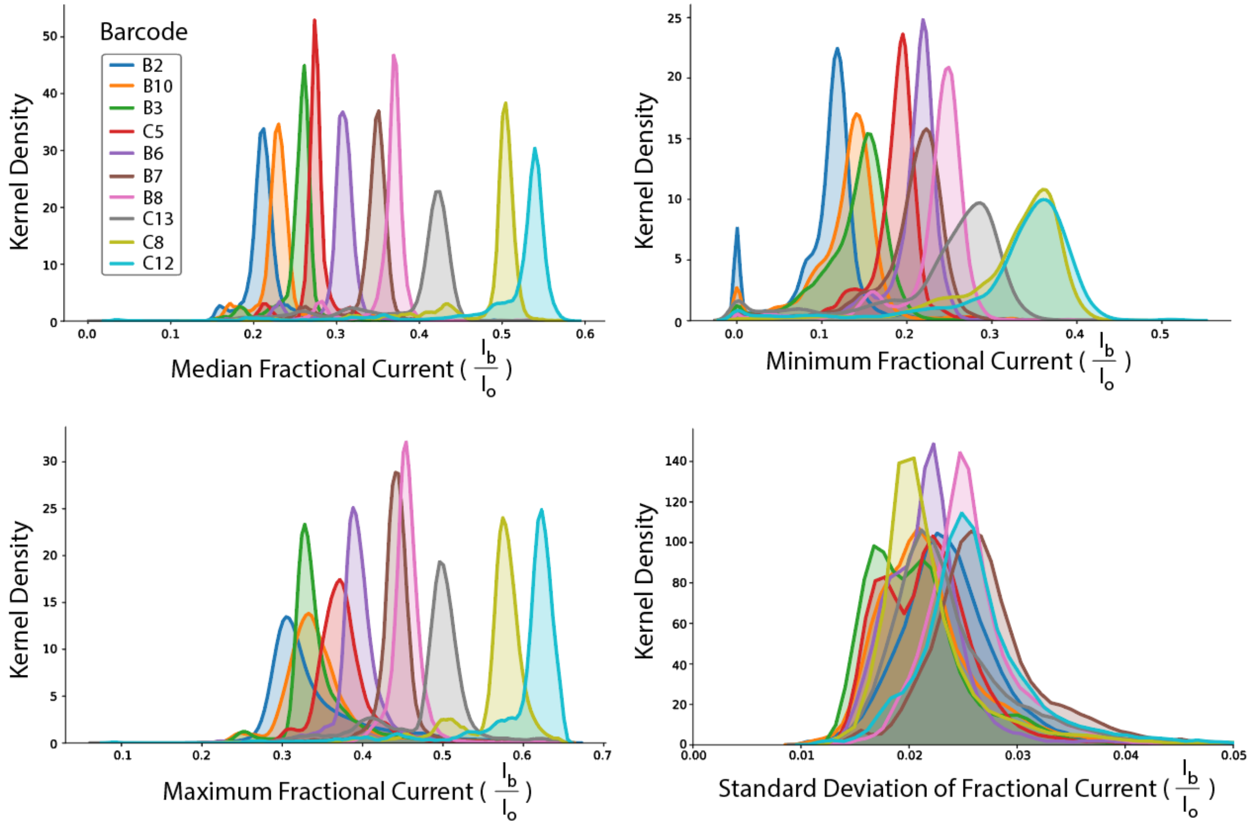


**Figure S5:** Distributions of median, minimum, maximum, and standard deviation of nanopore fractional current for each barcode in the selected set of orthogonal barcodes used for multiplexing experiments. Each distribution is composed of ~13000 data points.

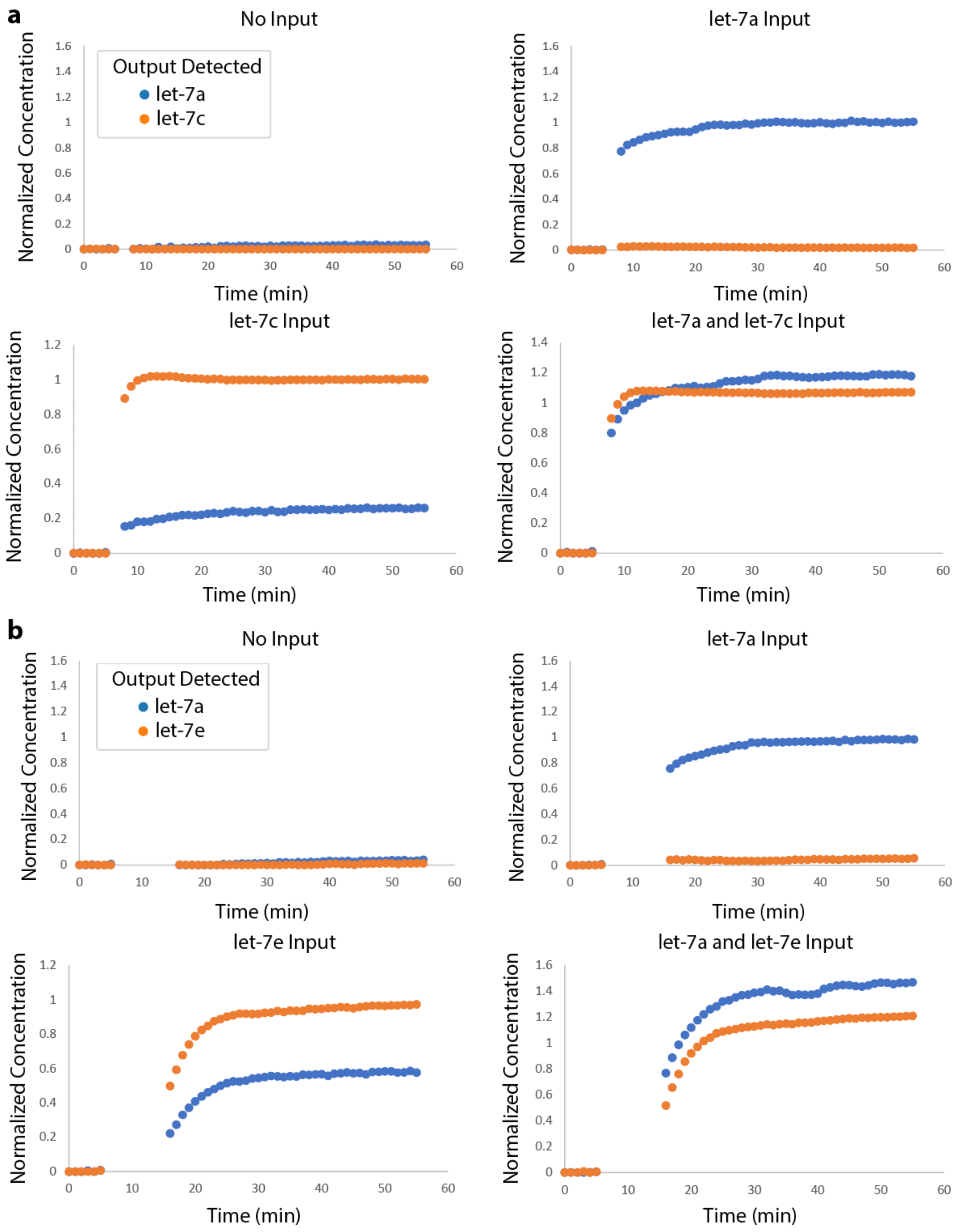
a
No Input
Output Detected
let-7a
let-7c
let-7a Input
let-7c Input
let-7a and let-7c Input
Normalized Concentration
Time (min)
b
No Input
Output Detected
let-7a
let-7e
let-7a Input
let-7e Input
let-7a and let-7e Input
Normalized Concentration
Time (min)

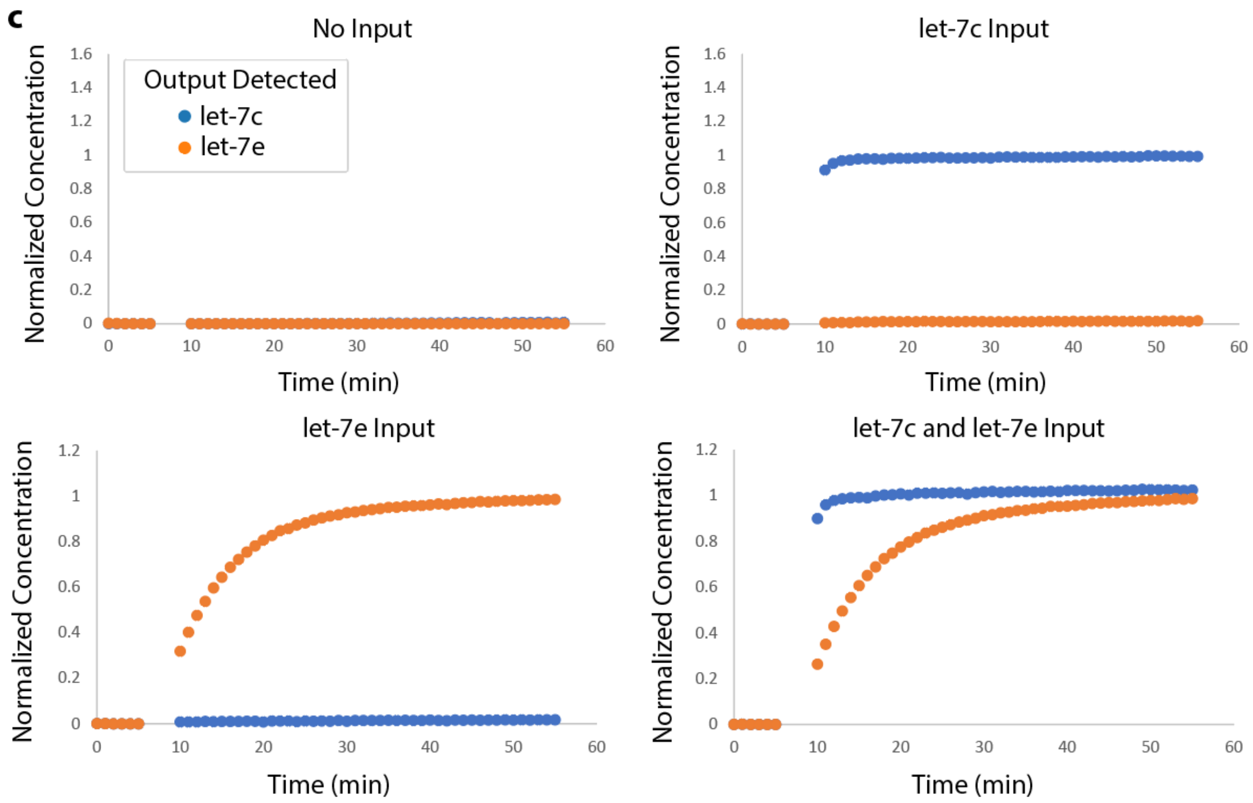


**Figure S6:** Normalized concentrations of let-7 probe output measured by fluorospectrometer. **a)** let-7a and let-7c probes, **b)** let-7a and let-7e probes, and **c)** let-7c and let-7e probes were multiplexed and their response to the introduction of miRNA input strands was measured. Each probe was present at 100 nM, each helper strand at 130 nM, each input strand at 50 nM, and streptavidin at 1200 nM.

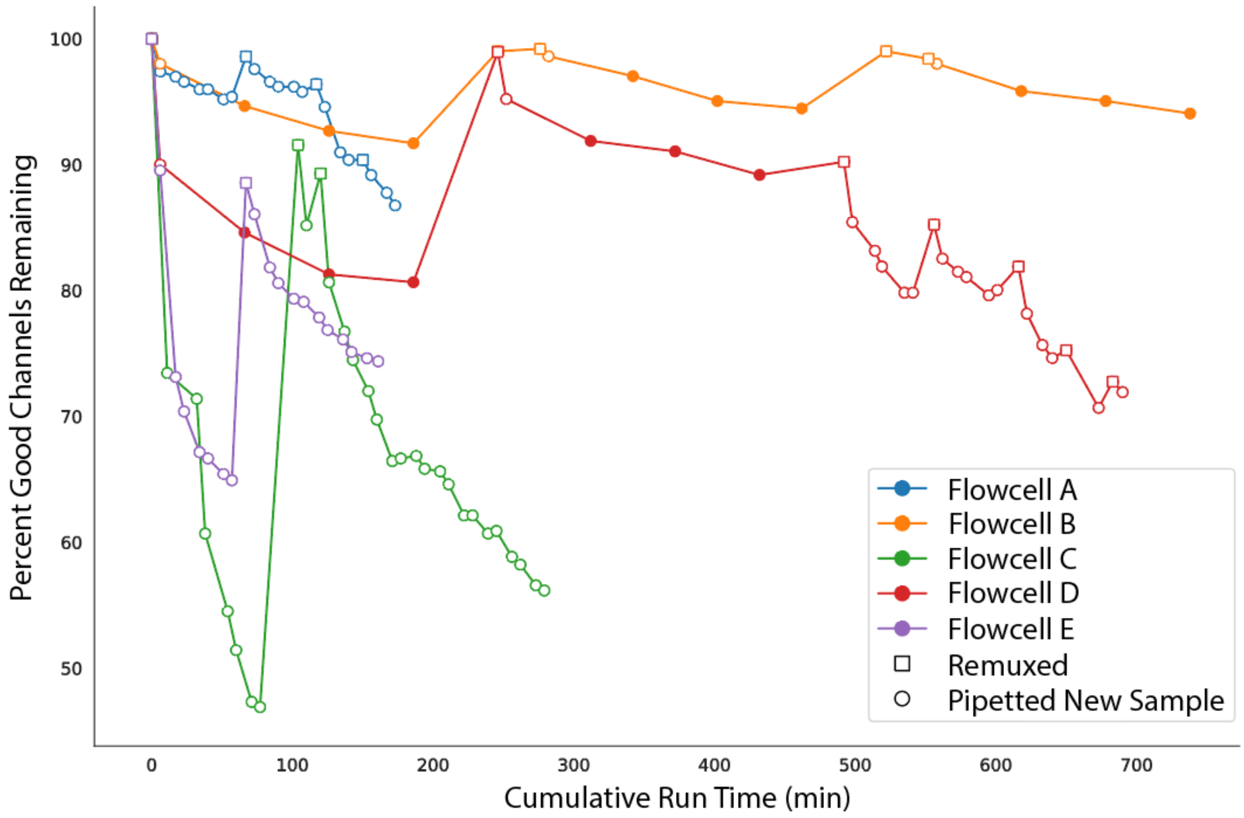


**Figure S7:** Plot showing the number of good channels on MinION R9.4.1 flow cells over run time. Each flow cell (colored line) begins with 400-512 good channels. Channel loss occurs at a higher rate when new samples (circles) are pipetted into the flow cell more frequently. Remuxing (squares) of the flow cell recovers channels.
